# Genome-wide identification and analysis of enhancer regulated microRNAs across 31 human cancers

**DOI:** 10.1101/2020.02.21.960351

**Authors:** Fei Tang, Mei Lang, Wenzhu Wang, Yanjing Li, Changying Li, Zheng tang Tan, Junjie Yue, Zhiyun Guo

## Abstract

Enhancers are cis-regulatory DNA elements that positively regulate the transcription of target genes in a tissue-specific manner and dysregulation in various diseases such as cancer. Recent studies showed that enhancers can regulate miRNAs and participate in the biological synthesis of miRNAs. However, the network of enhancer-regulated miRNAs across multiple cancers is still unclear. Here, a total of 2,418 proximal enhancer-miRNA interactions and 1,280 distal enhancer-miRNA interactions were identified through the integration of genomic distance, co-expression, and 3D genome data in 31 cancers. The results showed that both proximal and distal interactions exhibited significant tissue-specific feature and there was a noteworthy positive correlation between the expression of miRNA and the number of regulated enhancers in most tissues. Furthermore, it was found that there was a high correlation between the formation of enhancer-miRNA pairs and the expression of eRNAs whether in distal or proximal regulation. The characteristics analysis showed that miRes (enhancers that regulated miRNAs) and non-miRes presented significant differences in sequence conservation, GC content and histone modification signatures. Notably, GC content, H3K4me1, H3K36me3 were present differently between distal regulation and proximal regulation, suggesting they might participate in chromosome looping of enhancer-miRNA interactions. Finally, we introduced a case study, enhancer: chr1:1186391-1186507∼miR-200a was highly relevant to the survival of thyroid cancer patients and a cis-eQTL SNP on enhancer affected the expression TNFRSF18 gene as a tumor suppressor.

## Introduction

Enhancers are cis-regulatory DNA regulatory elements that positively regulate the transcription of target genes in a spatiotemporal-specific manner. The dysfunction of enhancer has been considered to affect enhancer-promoter communication and cause lots of diseases such as cancer (1). Previous studies have shown that enhancer activity is affected by the enhancer RNA (eRNA) which is transcribed bidirectionally from active regulatory enhancers, and plays a key role in regulating downstream gene expression. The Functional Annotation of the Mammalian Genome (FANTOM) group which applied CAGE technology had identified ∼ 65,000 active enhancers across multiple tissues, and these valuable resources provided important data sources for subsequent research (2). Recently, a large-scale pan-cancer study for TCGA patient samples across 33 cancer types revealed that the enhancer activity affects the expression of a variety of tumor-associated genes and was involved in tumor tumorigenesis (3). MicroRNAs (miRNAs) are a subset of endogenous non-coding RNAs (∼22 nucleotides long) which play vital roles in regulating genes expression via targeting the specific sites in 3’ untranslated region (3’ UTR) of miRNA (4, 5). In the past years, a great deal of literature confirmed that miRNAs are involved in almost all known cancers. A recent study showed that miR-24-1 which is present in nuclear promotes gene expression by targeting enhancers, suggesting there is an obvious interaction between enhancer and miRNA. Other recent studies showed that enhancers (including typical enhancers and super enhancers) are found to regulate miRNA expression and participate in the biological synthesis of miRNAs regulated by Drosha / DGCR8 (6, 7). These studies suggested that enhancers are involved in miRNA regulatory networks and contribute greatly to tumorigenesis and development.

However, the network of enhancer-regulated miRNAs across multiple tumors is still unclear. Therefore, a pan-cancer study was performed for enhancer-regulated miRNAs across 33 human cancer types in TCGA. Based on the distance between enhancer and miRNA, enhancer-miRNA pairs were classified into two types: proximal and distal enhancer-miRNA regulation. A series of enhancer-miRNA regulation were identified through the integration of co-expression, distance information, 3D genomes data of enhancers, and miRNAs from 8,693 samples in TCGA. GO and KEGG enrichment showed that target genes of enhancer-regulated miRNAs were significantly involved in tumor-associated biological processes and signaling pathways. Furthermore, it was found that there was a high correlation between the formation of enhancer-miRNA pairs and the expression of eRNAs. The results showed that miRes (enhancers that regulated miRNAs) and non-miRes presented significantly different characteristics including sequence conservation, GC content and histone modification signatures. Several histone modifications revealed significant cancer specificity and enhancer-miRNA spatial distance specificity. Finally, a case study was introduced, enhancer: chr1:1186391-1186507∼miR-200a was highly relevant to the survival of thyroid cancer patients and the cis-eQTL SNP on enhancer affected the expression of TNFRSF18 gene as a tumor suppressor.

## Materials and Methods

### 1.1 Identification of enhancer-miRNA interactions

Enhancer annotations and expressions data for TCGA 33 cancers were downloaded from a previous study (3). The expression data of miRNAs from all paired tumors and eight adjacent normal tissues were downloaded from the TCGA database. Co-expression analysis of enhancers and miRNAs was performed by using spearman correlation analysis (correlation coefficient |R| > 0.1, p-value<0.05).

Based on the distance of the enhancer-miRNA interactions, they could be classified into two types: proximal regulation and distal regulation. Referring to a previous study(8), proximal enhancer-miRNAs were calculated by the following formula: S (B/A) = (M−G) / (M+G). M and G represented the distance from the enhancer to the nearest miRNA gene and the nearest gene, respectively. The parameters A and B represent (G+M) / 2^1/2^ and (G-M) / 2^1/2^, respectively. S < 0.2 was adopted as the threshold to screen the reliable enhancer-miRNA pairs. Distal regulation of enhancer-miRNA was identified as the following procedure. Firstly, the transcription initiation sites (TSS) of 2,248 miRNAs were downloaded from the FANTOM5 data portal (9), 0.5 kb downstream and 1 kb upstream of the TSS of these miRNAs were defined as putative promoter region. A total of 1215 miRNAs were obtained through integrating 1881 miRNAs of TCGA and 2248 miRNAs of FANTOM5. Human chromatin interaction data were downloaded from 4DGenome (10). If the enhancer and miRNA promoter regions overlap with the chromatin interaction region of the 4D genome, it is considered that there is a physical interaction between the enhancer and miRNA, and the pair is defined as distal regulation.

### 1.2 Characteristics of enhancer-miRNA interactions

Enhancer RNAs (eRNAs) were determined by aligning the RNA transcribed from enhancer with the annotated RNA (GENCODE.v19). The transcripts overlapping protein-coding genes were removed. The GC content data were downloaded from the UCSC GC Percent track. The GC content was taken as the average of the regions of the enhancer itself. The PhastCons score was obtained from the UCSC cons100way track. The region of upstream and downstream which was 1 kb from the center of enhancer was considered as the calculation range of conservation.

The nine obtainable histone modification CHIP-Seq data of eight cell lines were downloaded from the ENCODE including H3K4me3, H3K4me1, H3K27ac, H3K9me3, H3K27me3, H3K36me3, H3K4me2, H3K9ac and H2K20me1. The eight cell lines matched eight types of cancer: A549 (LUAD), HepG2 (LIHC), HELA (CESC), HCT116 (COAD), DOHH2 (DLBC), PC-3 (PRAD), PANC-1 (PAAD), DND-41 (LAML). Signal consistency was considered when it appeared in at least five tissues.

### 1.3 Identification and analysis ubiquitously expressed enhancers

Enhancer-miRNA pairs that occurred in more than 10 tissues were defined as ubiquitously expressed enhancer-miRNA interaction. In order to investigate the function of the miRNA involved in enhancer-miRNA interaction, we downloaded the experimentally confirmed miRNA target genes from the miRTarbase database. Furthermore, target genes of each miRNA were subjected to GO and KEGG signaling pathway database for functional enrichment analysis using R package clusterProfiler (p.adjust <0.05). The eQTL data were retrieved from PancanQTL database (11), and a high correlation between SNP located on the enhancers and gene could be identified if the q value was lower than 0.05. Next, based on the database starBase(12), the expression level of the target miRNA inferred for the enhancer in the disease was analyzed by patient survival.

## Results and Discussion

### 2.1 Genome-wide identification of enhancer-miRNA interactions in 31 cancers

Previous studies have shown that enhancers are involved in the synthesis and regulation of miRNAs (13). To further explore the mechanism of enhancer-miRNA regulation in cancers, we identified a series of enhancer-regulated miRNAs in 33 cancer types. The co-expression between 15,080 enhancers from 8,693 samples and 1,881 miRNAs in 33 cancers was first analyzed. Finally, all co-expression pairs of enhancer-miRNA in 31 cancers were obtained except Uterine Corpus Endometrial Carcinoma (UCES) and Glioblastoma multiforme (GBM) because of too few enhancers and miRNA samples in these two tissues. Based on the distance between enhancer and miRNA, enhancer-miRNA pairs were divided into two types: proximal and distal enhancer-miRNA regulation. For proximal regulation, the method presented in the previous study was used to calculate enhancer-regulated neighbor miRNAs (8). For distal enhancer-miRNA regulation, the enhancer-miRNA interactions were identified by Hi-C data from 4Dgenome. Finally, a total of 2,418 proximal enhancer-miRNA pairs and 1,280 distal enhancer-miRNA pairs were obtained through the integration of genomic distance, co-expression and interaction analysis in 31 cancer (Figure 1A, 1B, Supplementary Table S1 and Table S2). To investigate whether these enhancer-miRNA interactions were tissue-specific or ubiquitously expressed, we counted the frequency of occurrence of these two types of interactions appearing in 31 cancers. The results revealed that both proximal and distal interactions exhibited significant tissue-specific feature, with only a few number of regulations (1.2% and 2.5% in proximal and distal interactions, respectively) ubiquitously expressed (presented in more than 10 cancers) (Figure 1C).

**Figure 1.**
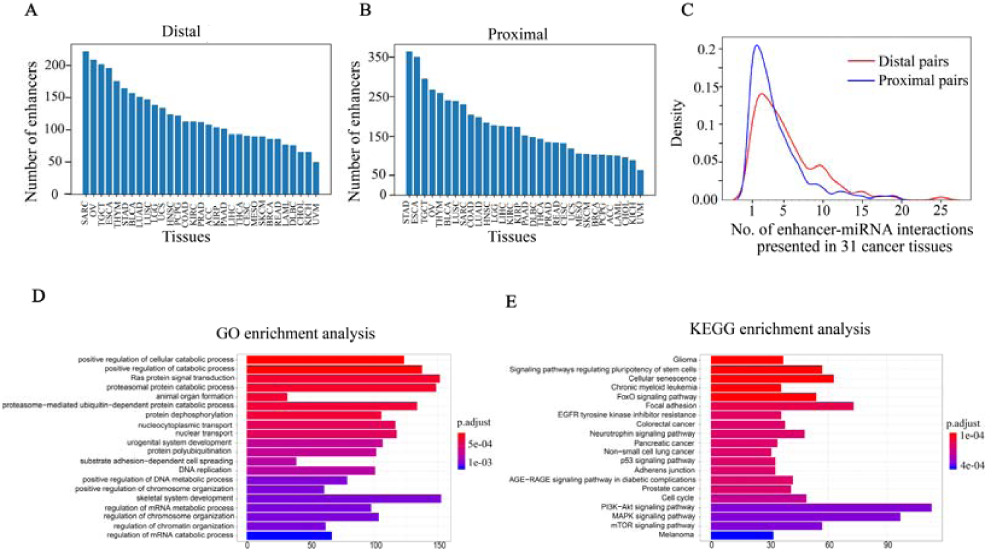
Go and KEGG pathway enrichment analysis of enhancer-miRNA regulation. No. of pairs of proximal regulations (A) or distal regulations (B) presented in each tissue. (C) The frequency of occurrence of enhancer-miRNA interactions appearing in 31 cancers. GO (D) and KEGG pathway (E) enrichment analysis of target genes of miRNAs that regulated by ubiquitously expressed enhancers.

If the regulatory relationship appears across a large number of cancers, it suggests that these regulations are critical for the tumorigenesis and development. To explore the biological functions of these miRNAs which are involved in ubiquitously expressed enhancer-miRNA regulation, the GO and KEGG functional enrichment analysis were performed using experimentally verified miRNA target genes. GO analysis indicated that these target genes of miRNAs that were regulated by ubiquitously expressed enhancers were significantly involved in tumor-associated biological processes such as cell cycle, cell differentiation, cell growth, metabolic regulation, metastasis, Ras protein catabolic process, etc. in distal (Figure 1D) or proximal regulation (Supplementary Figure S1A); KEGG analysis revealed that these miRNA target genes were significantly involved in cancer transcriptional dysregulation signaling pathways, such as FoxO signaling pathway, p53 signaling pathway, MAPK signaling pathway, the P13K-Akt signaling pathway, etc. in distal (Figure 1E) or proximal regulation (Supplementary Figure S1B).

### 2.2 The correlation between the miRNA expression and the number of regulated enhancers

Enhancers often regulate target genes and do not strictly follow one-to-one regulatory relationships. In order to investigate whether there is a correlation between the expression level of miRNAs and the number of miRes regulating these miRNAs, we performed principal component analysis (PCA) of the expression levels of miRNAs regulated by enhancers in 31 cancers. Here, only the distal regulation was analyzed because most enhancers-miRNA interactions in proximal regulation followed one-to-one regulatory rule according to genomic position restriction. The PCA results showed that the 31 cancers could be divided into three groups according to the number of the highly expressed miRNAs that were regulated by enhancers as follows: low group(1-3), medium(4-7) and high(>7) (Figure 2A, Supplementary Table S3). For example, miRNAs in PRAD, LUAD, LAML and ESCA regulated by more than seven enhancers presented significantly higher expression compared with the number of miRNAs regulated by enhancers that were less than seven (p<0.05). Interestingly, some similar types of cancers tended to cluster into one group which shared the same enhancer-miRNA regulation pattern. For example, the highest expression miRNAs in three types of kidney cancers (ACC, KIRC, KIRP) tended to be regulated by 4-7 enhancers (Figure 2A). Notably, there was a significant positive correlation between the expression of miRNA and the number of regulated enhancers in the Glad urothelial carcinoma (BLCA), Lung squamous cell carcinoma (LUSC), Ovarian serous cystadenocarcinoma (OV) and Testicular Germ Cell Tumors (TGCT) (Figure 2B, Supplementary Figure S2).

**Figure 2.**
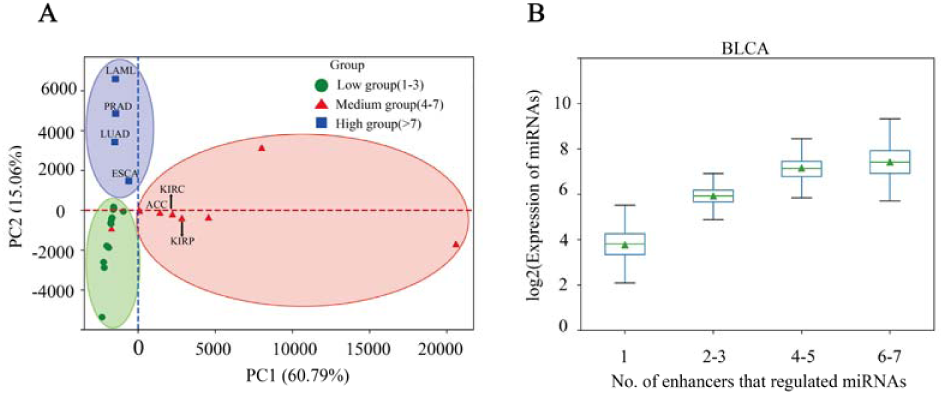
The correlation between miRNA expression and the number of regulated enhancers. (A) The PCA analysis of the expression levels of miRNAs regulated by enhancers in 31 cancers. (B) The expression of miRNA that regulated by different numbers of enhancers in BLCA.

### 2.3 There are significant differences in the sequence characteristics of miRes

It is important to explore the sequence characteristics of the miRes to conduct further identification of enhancer-miRNA interactions. Previously, it was reported that eRNA can be used as a trans-acting element to participate in the regulation of target genes (14). Consequently, the human GENCODE annotation was first used to investigate the transcript types of the distal and proximal regulatory miRes. It was found that 312 of the 998 (31.34%) enhancers that regulated distal miRNAs could transcribe eRNA, and the largest proportion (70.71%) of RNAs was lincRNA (Figure 3A). Similarly, the largest proportion of lincRNA was also found being present in enhancers that regulated proximal miRNAs (Supplementary Figure S3A). Moreover, we investigated whether there was a correlation between the formation of enhancer-miRNA pairs and the expression of eRNAs. The result showed that there was a high correlation between them in distal regulation (chi-square test, p-value<1.8e^-3^) and proximal regulation (chi-square test, p-value<10e^-5^), which suggested that enhancer might regulate the expression of miRNAs with the participation of eRNAs (Supplementary Table S4 and Table S5).

**Figure 3.**
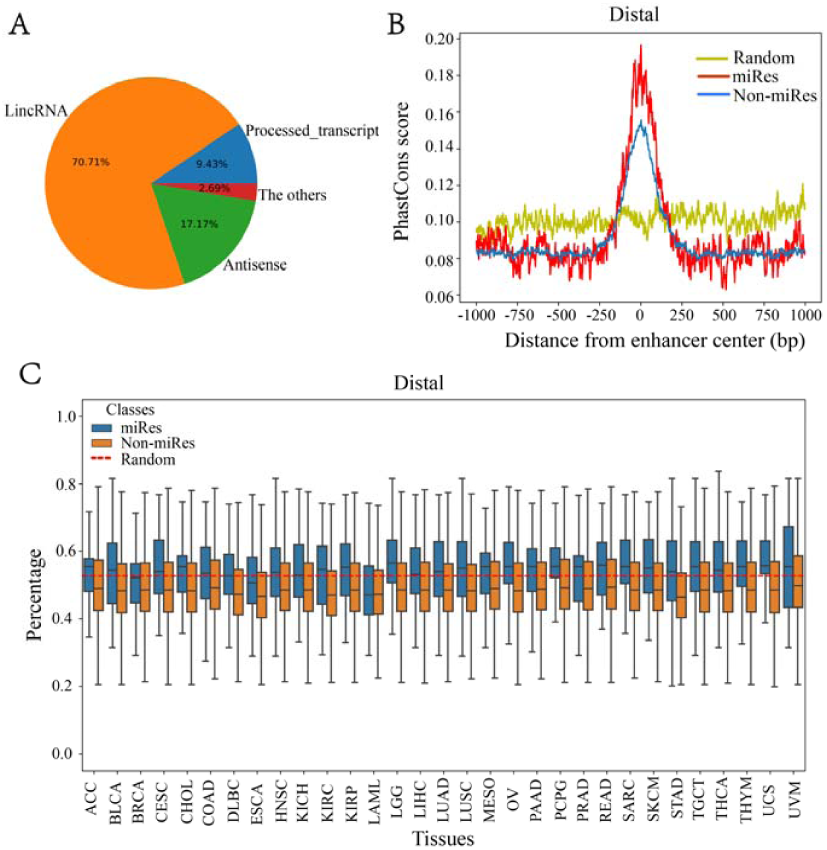
There are significant differences in the sequence characteristics of miRes. (A) Pie chart of all enhancer transcript in distal regulation. (B) Conservative score of the enhancer sequence using PhastCons in distal regulation. (C) GC content of the enhancer in distal regulation.

Continuously, PhastCons was used to analyze the conservation of the enhancer sequence. In distal regulatory, the results showed that the sequence of the enhancer was more conservative than the random sequence (p<3.2e^-23^), and the conserved region of enhancer was mainly located within ±250 bp around the center of enhancer (Figure 3B). Notably, the miRes showed higher conservation compared with the enhancers that did not regulate miRNAs. Similar results also appeared in proximal regulation (Supplementary Figure S3B). The above results indicated that the functional region of enhancer mainly concentrated near the enhancer center and that the miRes exhibited greater conservation than non-miRes. Furthermore, the GC content of the distal and proximal regulatory miRes were calculated. The results showed that miRes exhibited significantly higher GC content than the average value of random enhancer sequence in distal regulation(P-value<2.6e^-22^) and the miRe exhibited a higher GC content than non-miRe in each tissue (Figure 3C). Interestingly, there was no significant difference between the GC content of miRes and non-miRes in proximal regulation (P > 0.05) (Supplementary Figure S3C). Therefore, it was speculated that the GC content was an inherent property of the enhancer and might have a potential impact on chromosome looping which was more necessary in distal regulation than proximal regulation.

### 2.4 Histone modification showing cancer and miRes specific features

Previous studies have shown that the H3K27ac, H3K4me1 and H3K4me3 signal are key histone modification features for the activity of enhancers (15). To determine whether there are differences activity between miRes and non-miRes, we analyzed available H3K27ac, H3K4me1 and H3K4me3 ChIP-seq data in eight cancers. Not surprisingly, as an example shown in Figure 4A-F, all of the enhancers in distal regulation pairs and proximal regulation pairs had an enrichment of H3K27ac, H3K4me3 and H3K4me1 signal in the range of 1 kb upstream and downstream from the center of enhancer, and presented significantly higher signal in cancers than in normal tissues. Notably, the signals of H3K27ac and H3K4me3 of miRes were significantly higher than those of non-miRes in most tumor tissues (Supplementary Figure S4-7). Conversely, there was no significant difference in normal tissues. Interestingly, H3K4me1 showed that the difference between the miRe and non-miRe signals was only in distal regulation (Figure 4E, Supplementary Figure S8) but not in proximal regulation (Figure 4F, Supplementary Figure S9). This result was consistent with a previous study showing that enhancer activation of adjacent genes does not require H3K4me1 enrichment (16).

**Figure 4.**
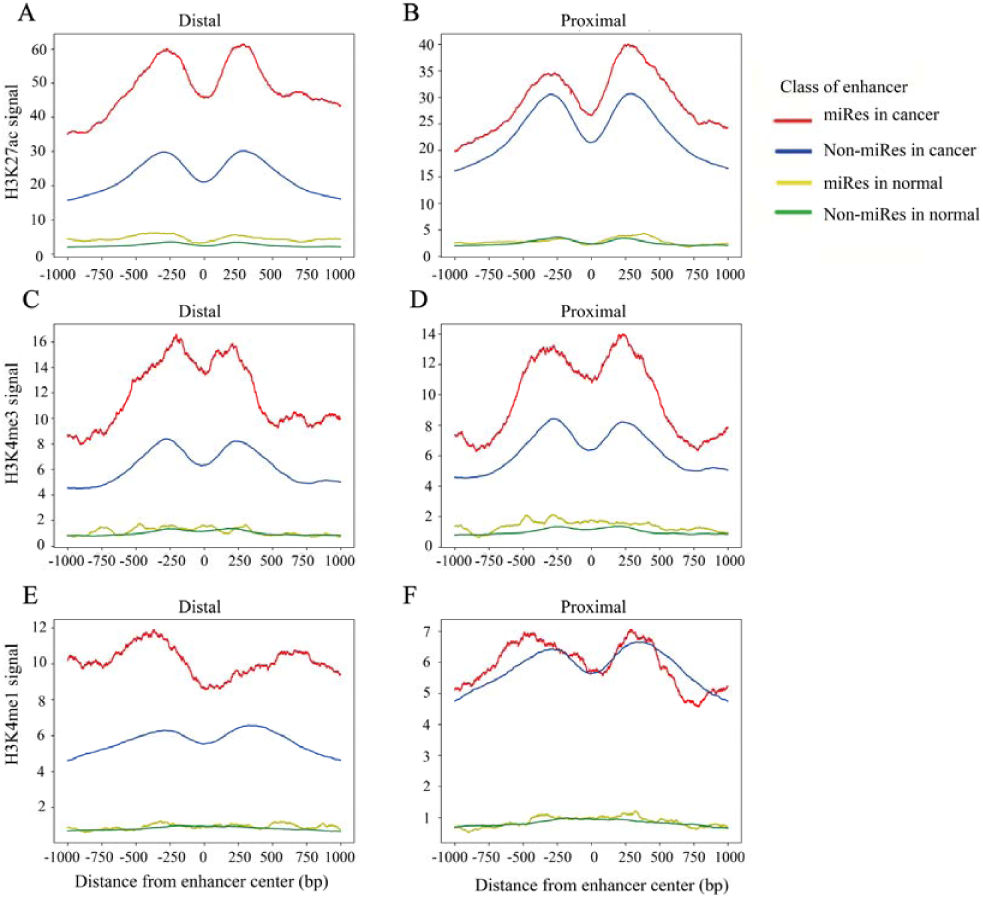
The signal of H3K27ac, H3K4me1 and H3k4me3 within ±1 kb surrounding the center of the enhancer center in LUAD.

In addition, it was asked if there were other histone modifications in addition to the above signals that had a significant difference between miRe and non-miRe. Therefore, we downloaded six histone modification data from ENCODE, including H3K4me2, H3K9ac, H3K20me1, H3K9me3, H3K27me3 and H3K36me3. It was found that H3K9me3 and H3K36me3 in distal and proximal pairs were significantly different between the miRes and non-miRes in at least five cancer tissues (Supplementary Figure S10-13). Among them, H3K9me3 showed lower enrichment in miRes compared with in non-miRes probably due to this histone modification which was the marker of heterochromatin (Supplementary Figure S10-11) (17). This result was consistent with our previous supposition that the transcription of enhancers had a positive effect on the expression of miRNA that enhancer regulated. H3K36me3, a marker for transcription extension, showed a high enrichment in the miRes in distal interaction pairs but not in the proximal interaction pairs. According to a previous study (18), transcriptional elongation has an effect on the spatial structure of chromatin, and this may have more influence on distal regulation than proximal regulation (Supplementary Figure S12-13).

### 2.5 A case study of miRe in thyroid carcinoma

To investigate miRes identified as described above, here we introduced a case study about an enhancer: chr1:1186391-1186507 and its target miRNA: miR-200a in thyroid carcinoma (THCA). A cis-eQTL SNP (rs6603785) identified on enhancer: chr1:1186391-1186507 is located close to the transcription start site (TSS) of TNFRSF18 gene, which acts as a tumor suppressor (19), and mainly occurs when the base A mutates to T (Figure 5A). There was a significant difference in the expression levels of samples of different genotypes (p<1.76e^-4^) (Supplementary Figure S14). In addition, miR-200a as a target of enhancer was highly relevant to the survival of thyroid cancer patients (Figure 5B). A previous study showed that miR-200 regulates the epithelial stromal transformation of thyroid cancer through EGF/EGFR signal (20) and it is a key factor in the epithelial phenotype and a tumor suppressor in thyroid carcinoma (21). Additionally, the survival analysis showed that low expression of miR-200a patients had a lower survival time.

**Figure 5.**
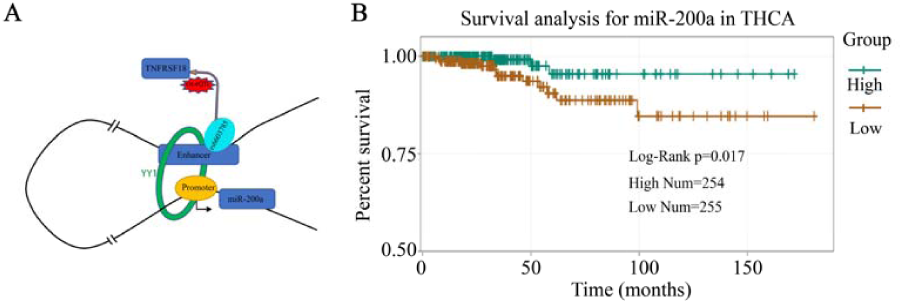
A case study of miRes in thyroid carcinoma. (A) An example of enhancer-miRNA regulation interaction (chr1:1186391-1186507∼miR-200a). (B) The survival curve of hsa-miR-200a in thyroid cancer.

## Acknowledgments

This work was supported by the National Science and Technology Major Project of Infectious Diseases [2018ZX10101-003-001-008] and National Natural Science Foundation of China (31671363).

