## Supplementary Figures for "Genome-wide identification and analysis of enhancer regulated microRNAs across 31 human cancers"

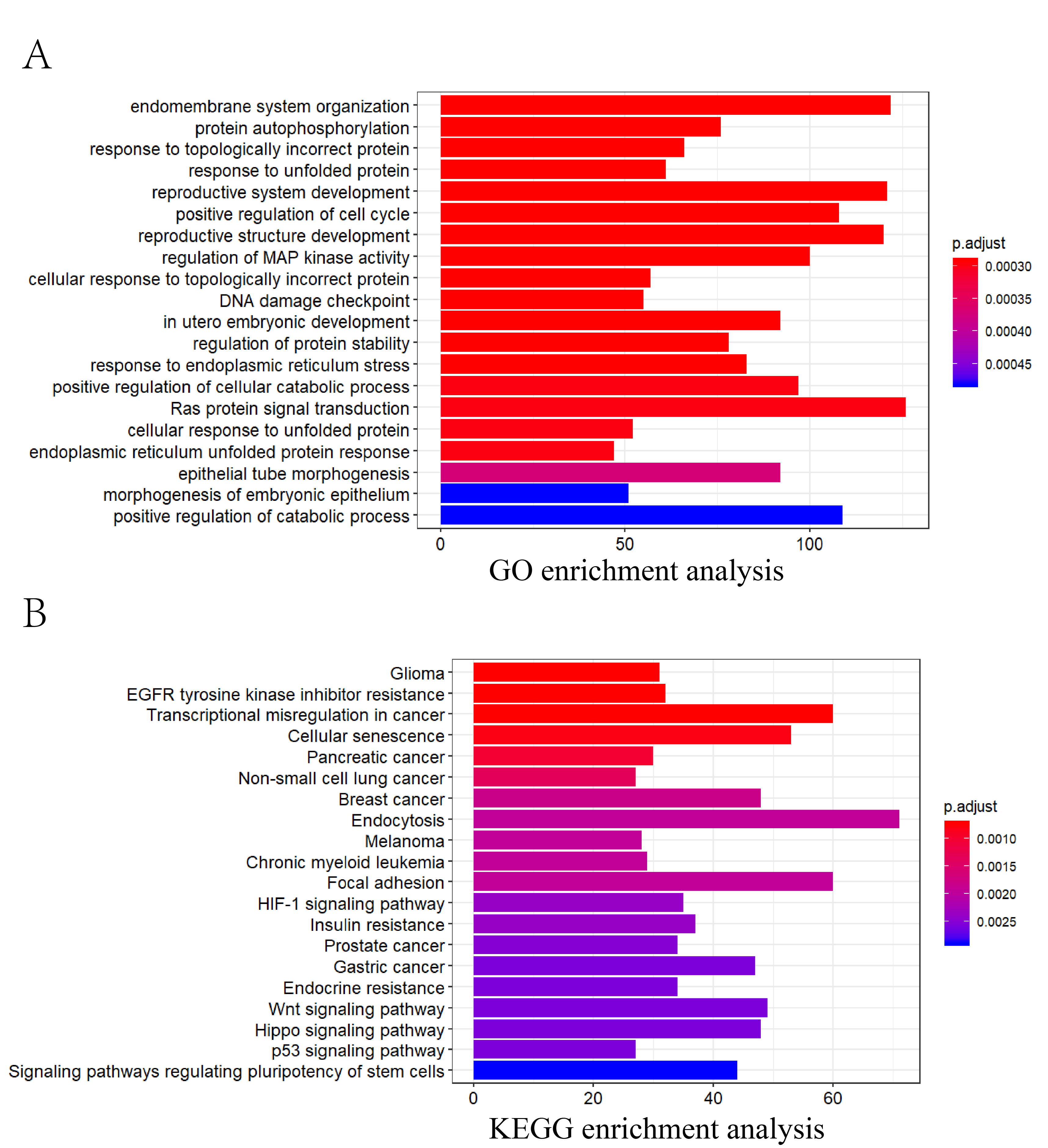

**Figure S1: Pathway enrichment analysis of proximal enhancer-miRNA regulation pairs. (A)** GO enrichment analysis of target genes of miRNAs that were regulated by ubiquitously expressed enhancers in proximal regulation pairs. Same as distal regulation pairs, GO analysis indicated that these target genes of miRNAs that were regulated by ubiquitously expressed enhancers were significantly involved in tumor-associated biological processes. (**B)** KEGG enrichment analysis of target genes of miRNAs that were regulated by ubiquitously expressed enhancers in proximal regulation pairs. Same as distal regulation pairs, these target genes of miRNA in proximal regulation pairs were significantly involved in cancer transcriptional dysregulation signaling pathways.

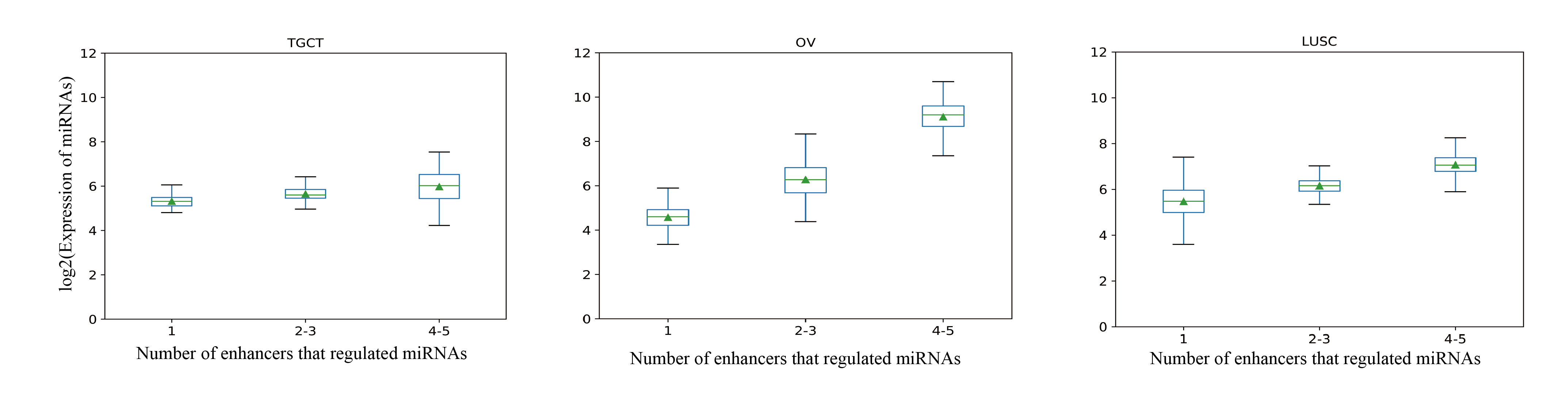

**Figure S2: Correlation between number of contraction enhancers and miRNA expression.** There was a significant positive correlation between the expression of miRNA and the number of regulated enhancers in the Testicular Germ Cell Tumors (TGCT), Ovarian serous cystadenocarcinoma (OV) and Lung squamous cell carcinoma (LUSC)

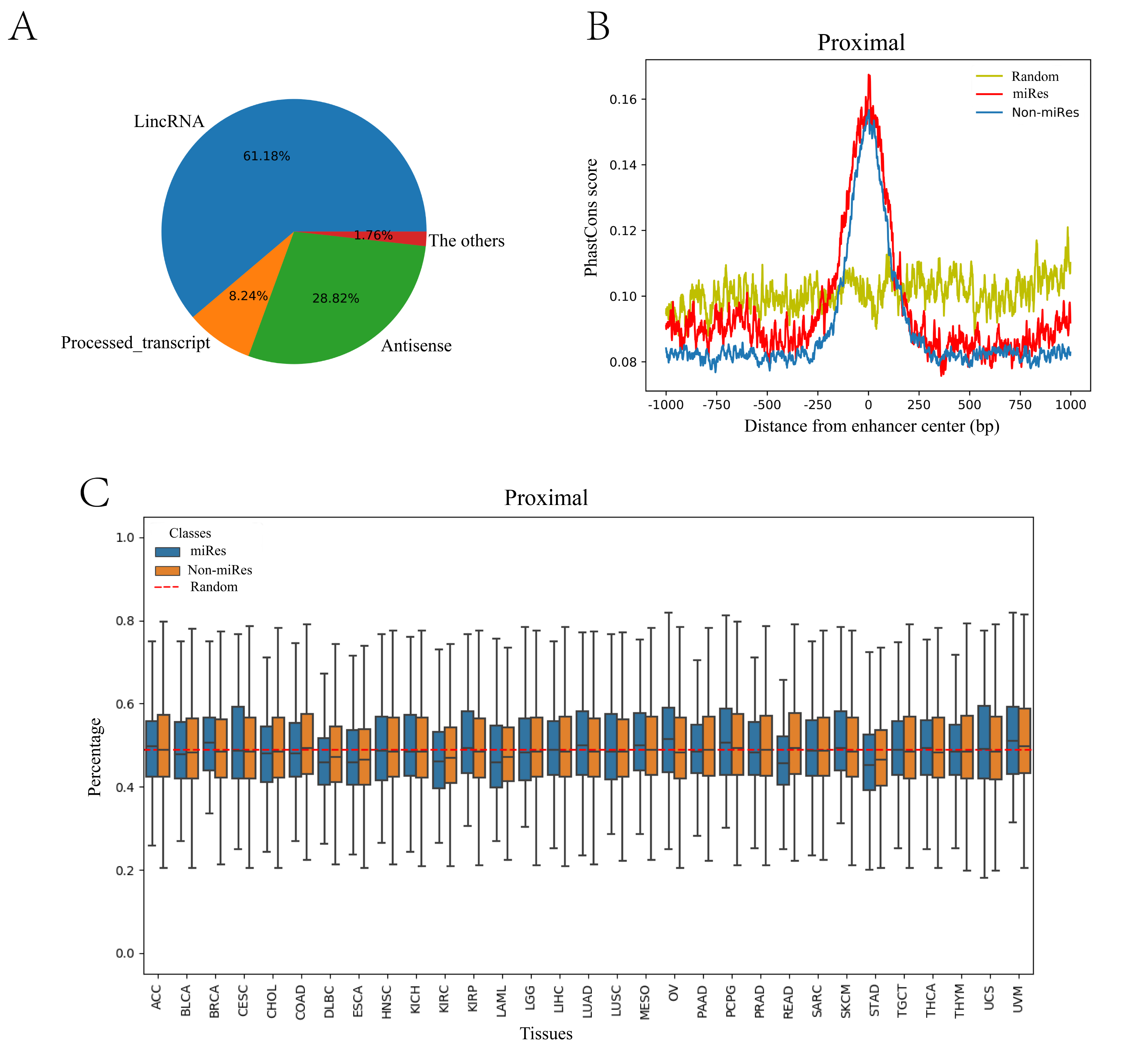

**Figure S3: Sequence characteristics of miRes and non-miRes in proximal regulation. (A)** Pie chart of all enhancer transcriptions in proximal regulation. 973 of the 2418 (31.34%) enhancers that regulated distal miRNAs could transcribe eRNA, and the largest proportion (61.18%) of RNAs was lincRNA. (**B)** Conservative score of enhancers in proximal regulation. The results showed that the sequence of the enhancer was more conservative than the random sequence (p<7.0e-28), and the miRes showed a higher conservation compared with the enhancers that did not regulate miRNAs. (**C)** GC content of the enhancer in proximal regulation. Unlike distal regulation pairs, there was no significant difference between the GC content of miRes and non-miRes in proximal regulation (P > 0.05).

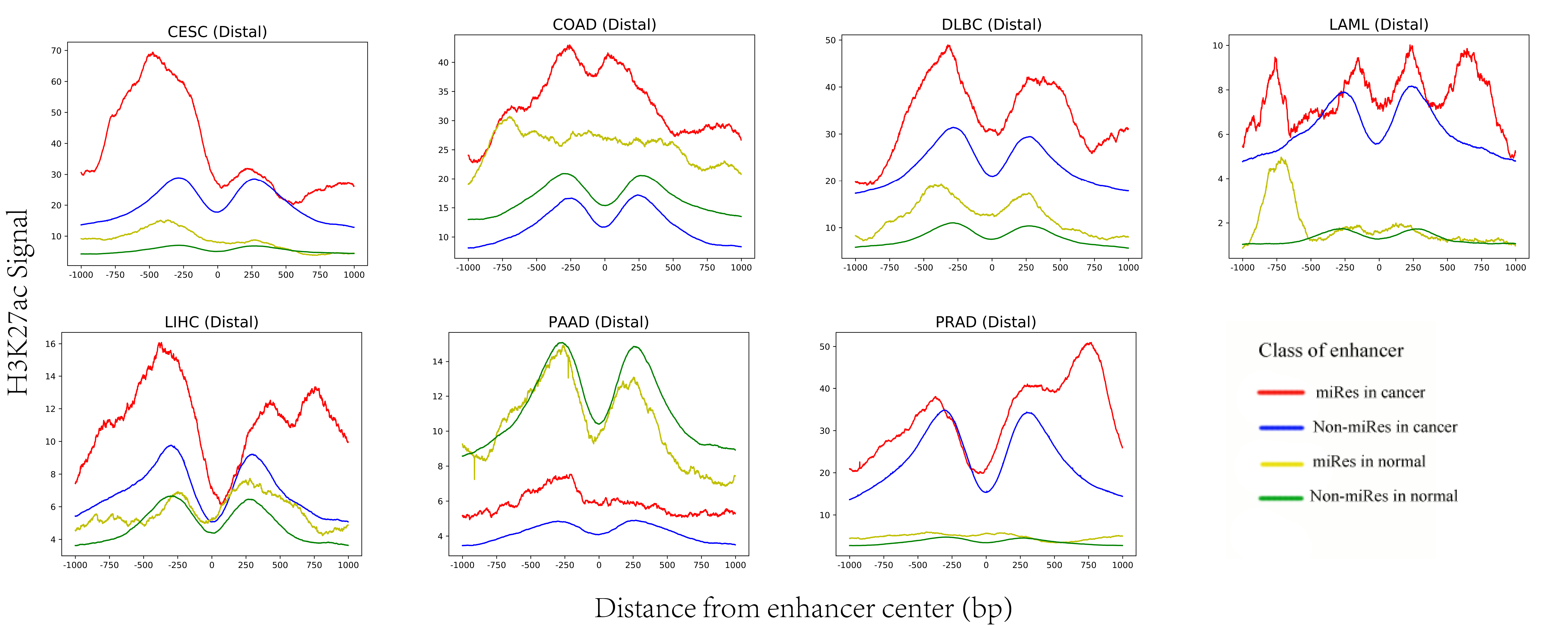
**Figure S4: The signal of H3K27ac within ±1 kb of the enhancer center in distal regulation pairs.**

The signals of H3K27ac of miRes in distal regulation pairs were significantly higher than those of non-miRes in most tumor tissues. There was no significant difference in normal tissues. Except for PAAD, the signal of enhancer in cancer tissue was higher than that of enhancer in normal tissue in most diseases.

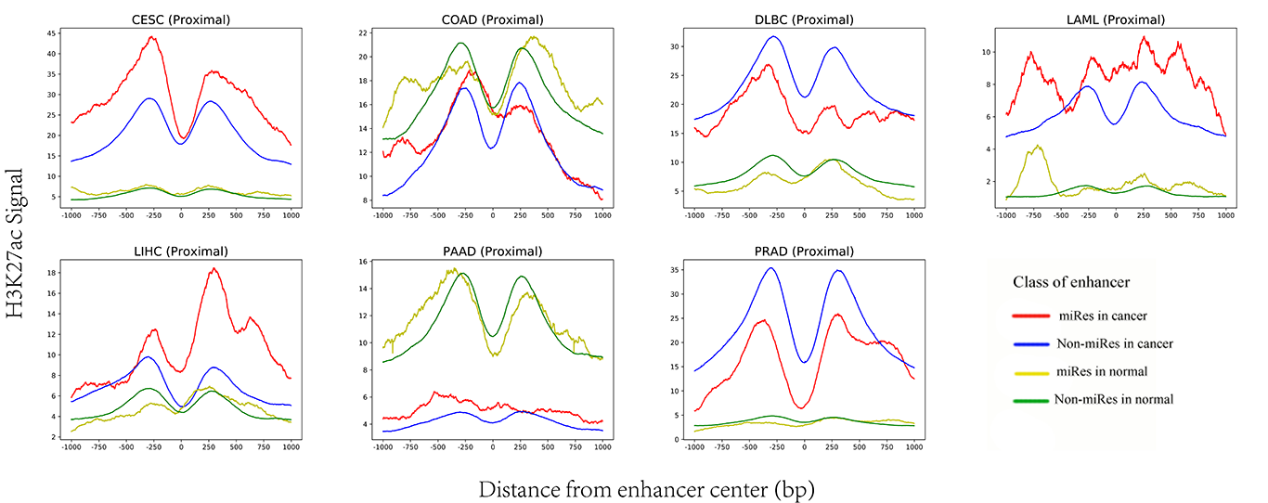

**Figure S5: The signal of H3K27ac within ±1 kb of the enhancer center in proximal regulation pairs.** The signals of H3K27ac of miRes in proximal regulation pairs were significantly higher than those of non-miRes in most tumor tissues. There was no significant difference in normal tissues. Except for COAD and PAAD, the signal of enhancer in cancer tissue was higher than that of enhancer in normal tissue in most diseases.

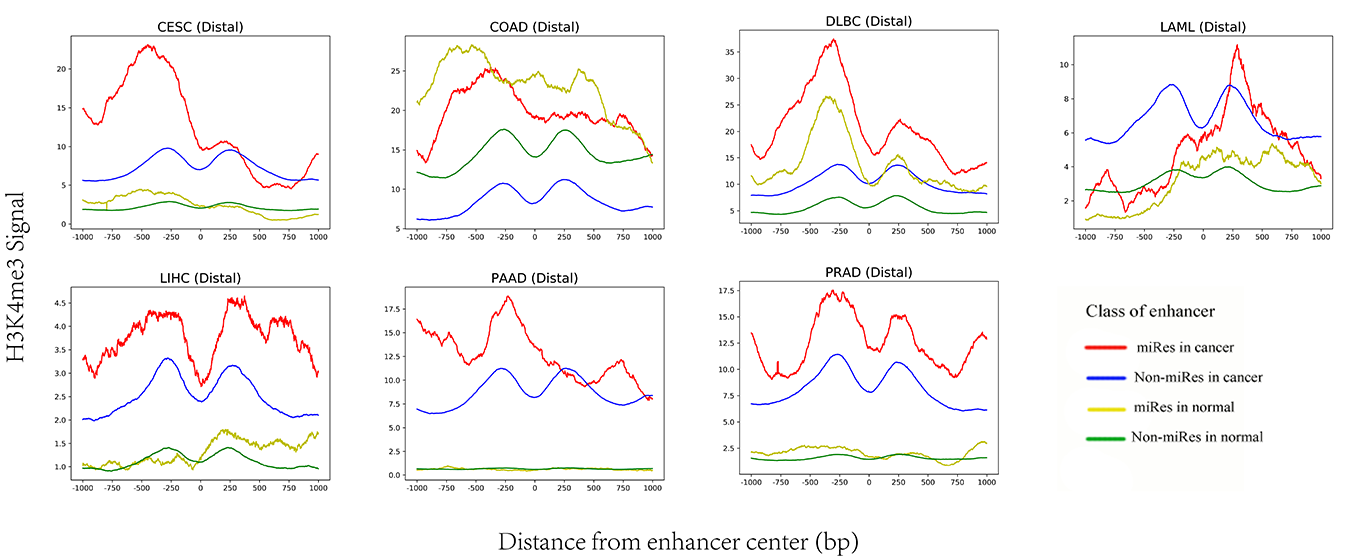

**Figure S6: The signal of H3K4me3 within ±1 kb of the enhancer center in distal regulation pairs.**

The signals of H3K4me3 of miRes in distal regulation pairs were significantly higher than those of non-miRes in most tumor tissues, and the signal of enhancer in cancer tissue was higher than that of enhancer in normal tissue in most diseases.

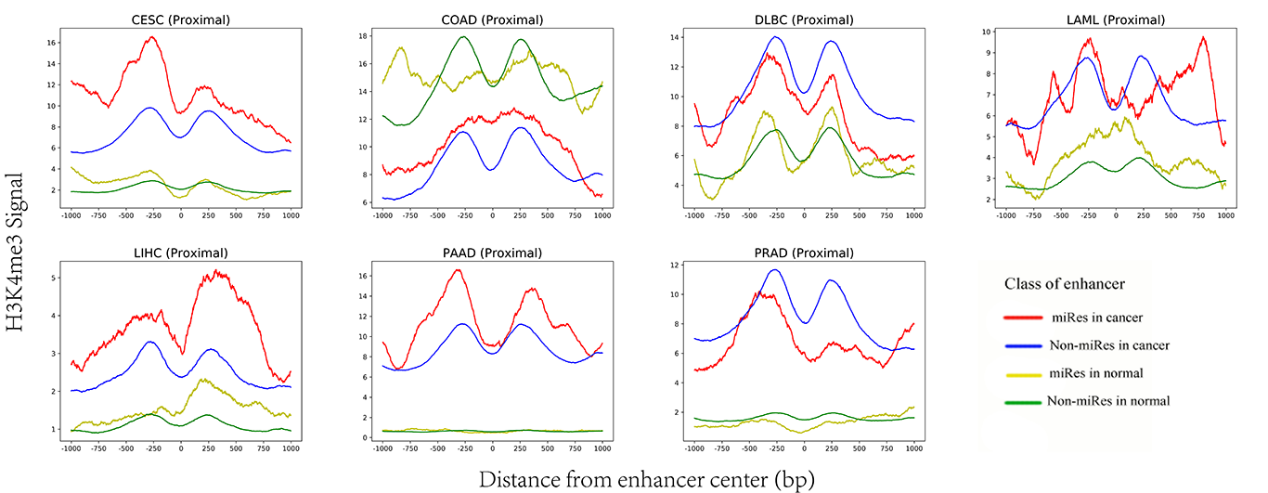

**Figure S7: The signal of H3K4me3 within ±1 kb of the enhancer center in proximal regulation pairs.** The signals of H3K4me3 of miRes in proximal regulation pairs were significantly higher than those of non-miRes in most tumor tissues. There was no significant difference in normal tissues. Except for COAD, the signal of enhancer in cancer tissue was higher than that of enhancer in normal tissue in most diseases.

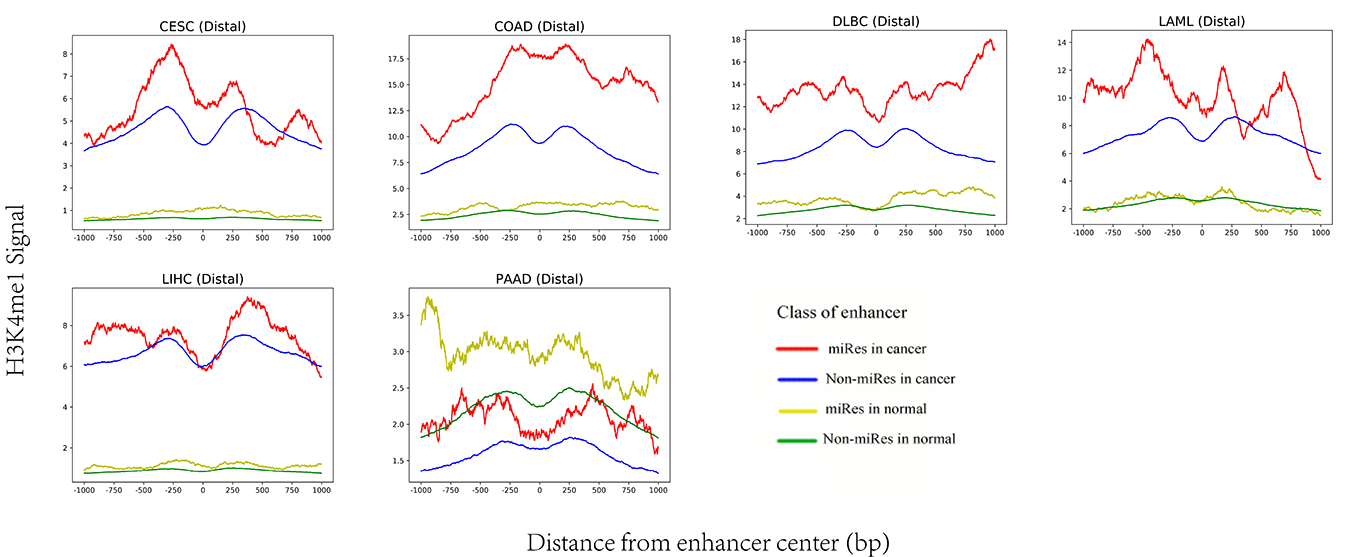

**Figure S8: The signal of H3K4me1 within ±1 kb of the enhancer center in distal regulation pairs.** The signals of H3K4me1 of miRes in distal regulation pairs were significantly higher than those of non-miRes in most tumor tissues. There was no significant difference in normal tissues. Expect for PAAD, the signal of enhancer in cancer tissue was higher than that of enhancer in normal tissue in most diseases.

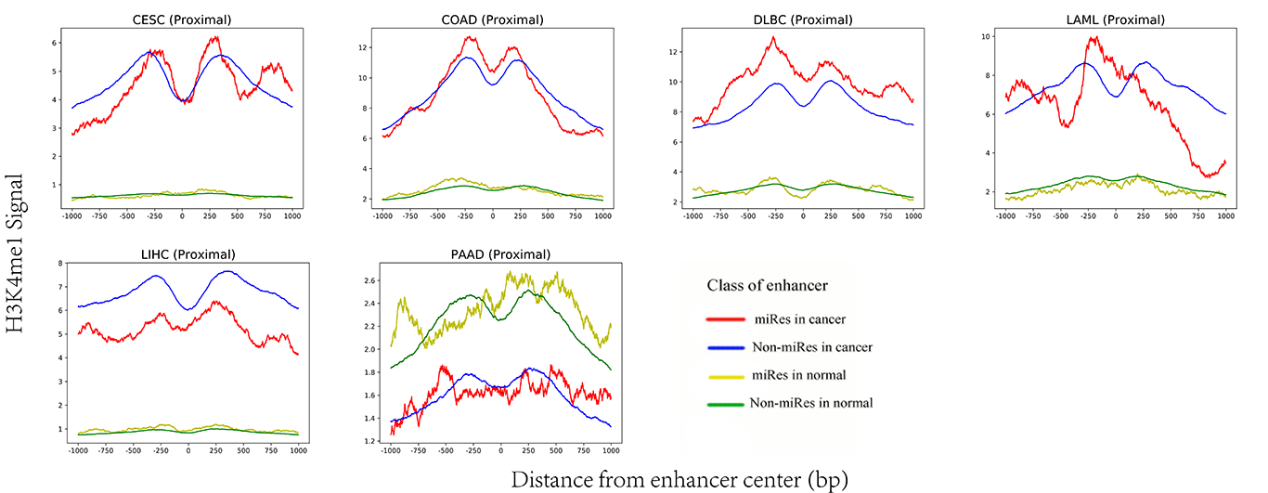

**Figure S9: The signal of H3K4me1 within ±1 kb of the enhancer center in proximal regulation pairs.** There was no significant difference between the H3K4me3 signal of miRes in proximal regulation pairs and that of non-miRes in most tumor tissues. There was no significant difference in normal tissues. Except for PAAD, the signal of enhancer in cancer tissue was higher than that of enhancer in normal tissue in most diseases.

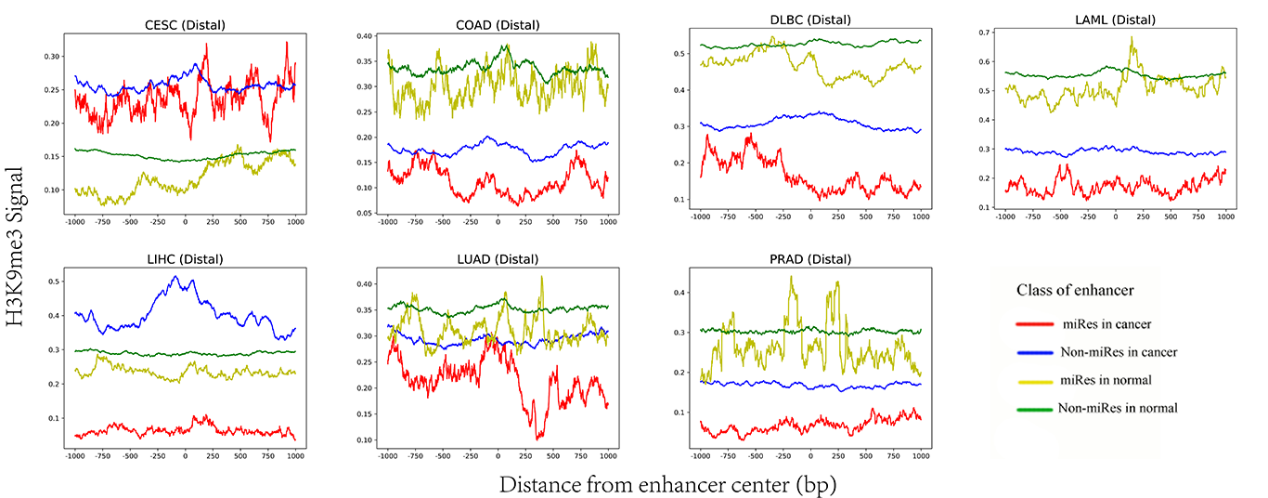

**Figure S10: The signal of H3K9me3 within ±1 kb of the enhancer center in distal regulation pairs.** The signal of H3K9me3 showed lower enrichment in miRes of distal regulation pairs compared with in non-miRes in most tumor tissues. There was no significant difference in normal tissues.

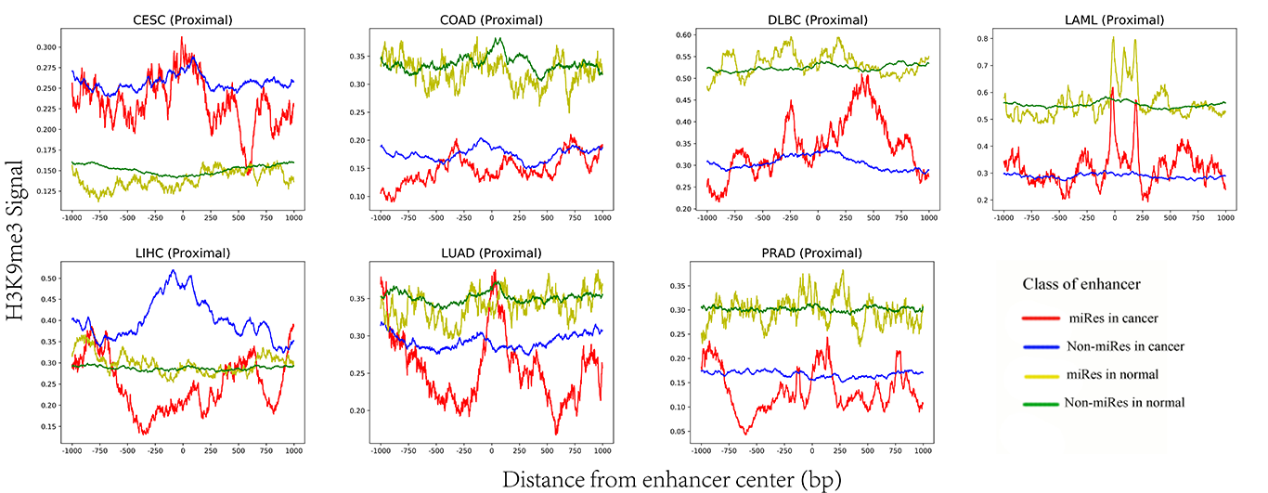

**Figure S11: The signal of H3K9me3 within ±1 kb of the enhancer center in proximal regulation pairs.** The signal of H3K9me3 showed lower enrichment in miRes of proximal regulation pairs compared with in non-miRes in most tumor tissues. There was no significant difference in normal tissues.

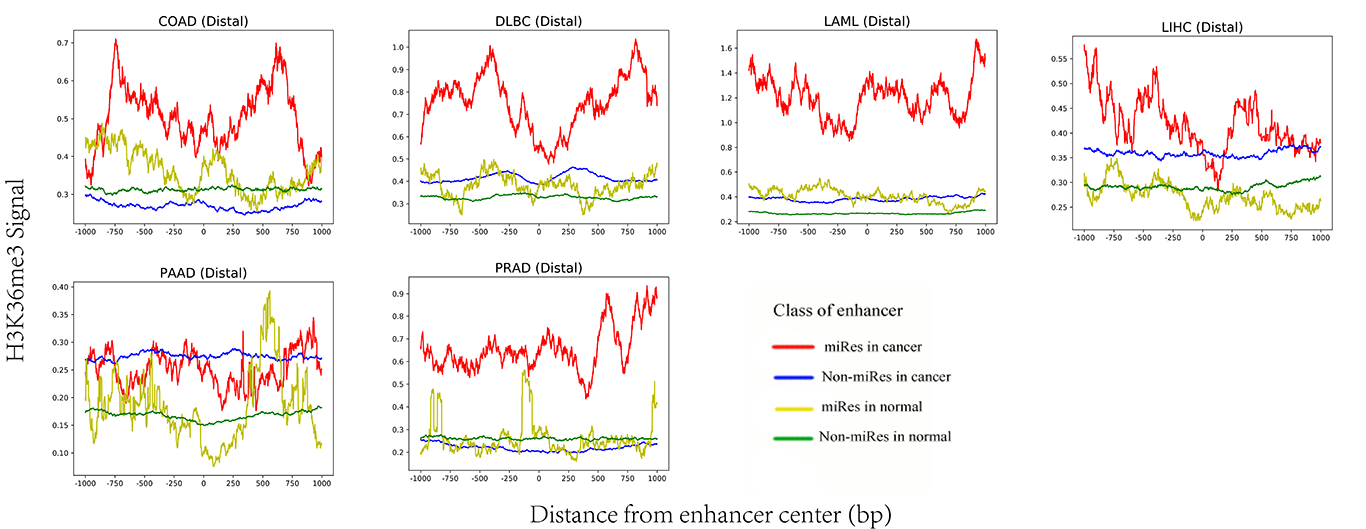

**Figure S12: The signal of H3K36me3 within ±1 kb of the enhancer center in distal regulation pairs.** The signal of H3K36me3 showed higher enrichment in miRes of distal regulation pairs compared with in non-miRes in most tumor tissues. There was no significant difference in normal tissues.

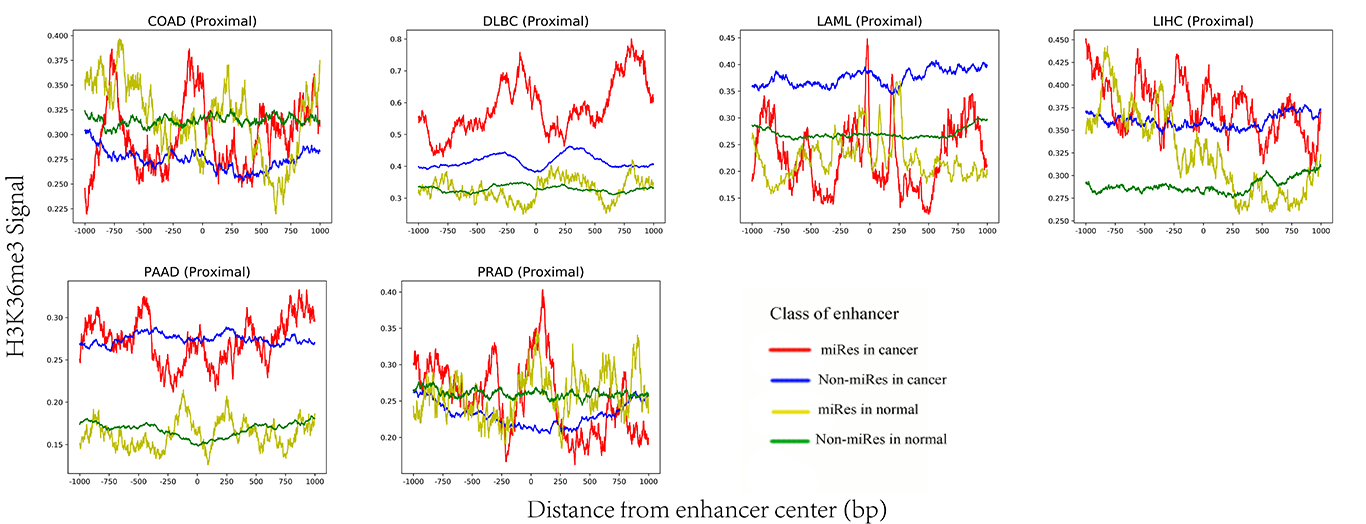

**Figure S13: The signal of H3K36me3 within ±1 kb of the enhancer center in proximal regulation pairs.** The signal of H3k36me3 had no significant difference in the miRe of distal regulation pairs in most tumor and normal tissues compared with that of non-miRe.

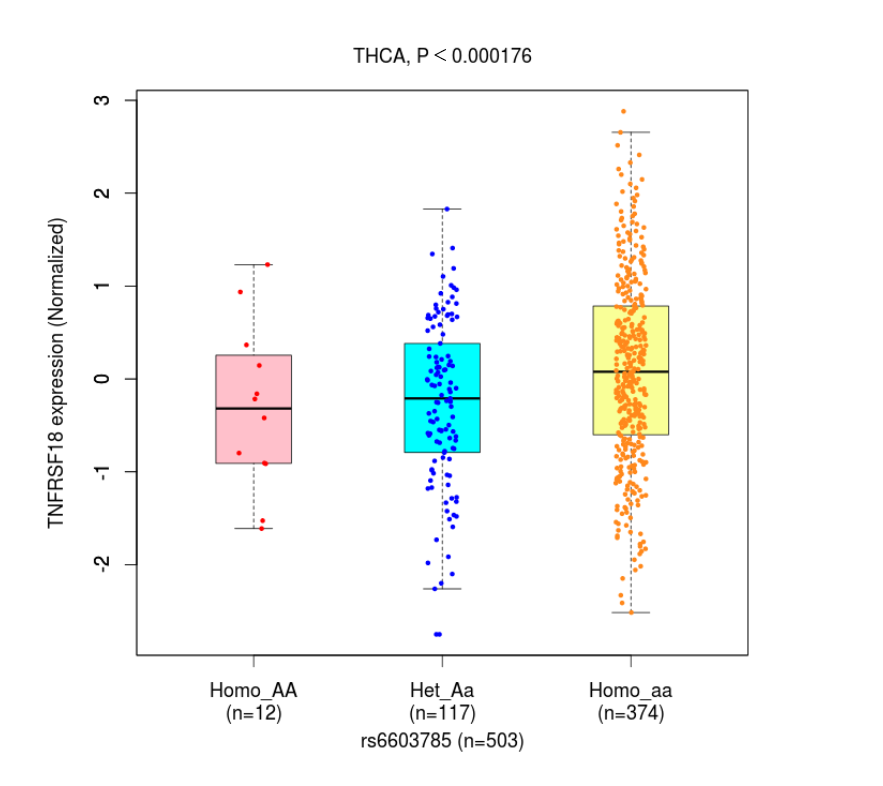

**Figure S14: Expression of three genotypes of rs6603785.** Significant differences in the expression levels of the three genotypes of this SNP (rs6603785).
