## Supplementary Table S3 for "Genome-wide identification and analysis of enhancer regulated microRNAs across 31 human cancers"

**Table S3.** PCA of the expression levels of miRNAs regulated by enhancers in 31 tissues

| Group | Disease |
|---|---|
| High group（>7） | ESCA，LAML，LUAD，PRAD |
| Medium group（4-5） | ACC, BRCA, COAD, KIRC, KIRP, PAAD, SARC |
| Low group（1-3） | CESC，CHOL，DLBC，HNSC，KICH，LGG，LIHC，MESO，PCPG，READ，SKCM，STAD，THCA，THYM，UCS，UVM，BLCA，LUSC，TGCT，OV |

**Ps:** Red diseases indicate the group with positive correlation.
