## Supplementary Table S4 for "Genome-wide identification and analysis of enhancer regulated microRNAs across 31 human cancers"

**Table S4.** Enhancers regulating miRNA associated with transcription of enhancers in distal regulation

| Number | miRes | Non-miRes | Total |
|---|---|---|---|
| Enhancer with transcript | 321 | 4109 | 4430 |
| Enhancer without transcript | 686 | 10692 | 11378 |
| Total | 1007 | 14801 | 15808 |

Chi square-value >7.71 P-value $<5.48e^{-4}$
