## Supplementary Table S5 for "Genome-wide identification and analysis of enhancer regulated microRNAs across 31 human cancers"

**Table S5.** Enhancers regulating miRNA associated with transcription of enhancers in proximal regulation

| Number | miRes | Non-miRes | Total |
|---|---|---|---|
| Enhancer with transcript | 973 | 3448 | 4421 |
| Enhancer without transcript | 1445 | 9942 | 11387 |
| Total | 2418 | 13390 | 15808 |

Chi square-value >212.72 P-value <$3.5e^{-47}$
